# CXADR is a Human IgG Fc Receptor

**DOI:** 10.1101/2024.10.22.619713

**Authors:** Christopher J. Emig, Adela A. Vitug

## Abstract

CXADR is a membrane protein present in epithelial tight junctions and other tissues with a variety of known functions, in addition to its role as entry receptor for coxsackie B virus and adenovirus 2 and 5. We identified a previously unknown function of CXADR as a binding partner for human IgG but not IgA, IgE, IgM, or ScFv in a non paratope specific manner. Binding was specific to human and rabbit IgG, and not observed for mouse, rat, donkey, or goat IgG. Binding of Ig to CXADR is inhibited by FcBlock (BD), and is competitive with anti-Fc binding secondary antibodies, but not anti-Fab secondary antibodies, indicating that CXADR binds the Fc portion of IgG. CXADR, being a member of the immune superfamily of proteins, has similar structural homology to other Ig binding proteins, including Fc receptors. We conclude that CXADR has a previously unknown function as a human Fc receptor, serving as a direct link between the humoral adaptive immune system and many cell types in the human body previously unknown to express FcR.

## Introduction

CXADR is a membrane protein present in epithelial tight junctions^1,2^ that plays a role in transepithelial migration of lymphocytes^3^, neurite outgrowth and synaptic function ^4,5^, activation of gamma-delta T cells ^6^, spermatogenesis^7^ and cardiomyocyte function^8–11^, in addition to its role as entry receptor for coxsackie B virus and adenovirus 2 and 5^12,13^. Mutations in or near CXADR are associated with cardiac arrythmias^9^ and COVID-19 induced cardiomyositis (Hacimustafaoglu et al., 2021), and Type I diabetes^14^. During a high throughput serology study, we observed an unexpected result in which all 135 patient plasma samples appeared to be seropositive to CXADR^15^. In this work we deconvolved the binding partners present in human plasma responsible for CXADR binding and characterized the selectivity, affinity and epitope specificity of the primary binding partner identified, human IgG.

## Results

### Binding to human plasma

Every human plasma sample demonstrated saturated binding to CXADR at dilutions of 1:500, 1:1000 (Figure 1) and 1:2000, while secondary antibody had no detectable signal with a PBS only condition. Plasma samples were also measured for binding to HSV-1 Lysate, which demonstrated a diversity in signal expected for a heterogeneous population of unknown serostatus. Human plasma was diluted 1:1000 in PBS and assayed for CXADR binding using an anti-Fab specific to human IgA, IgM and IgG. CONSV3 control serum (Sigma-Aldrich, Burlington, MA) was included as a positive control for HSV1-Lysate (Native Antigen Company, Oxford, UK) binding and PBS only as a negative control.

**Figure 1.**
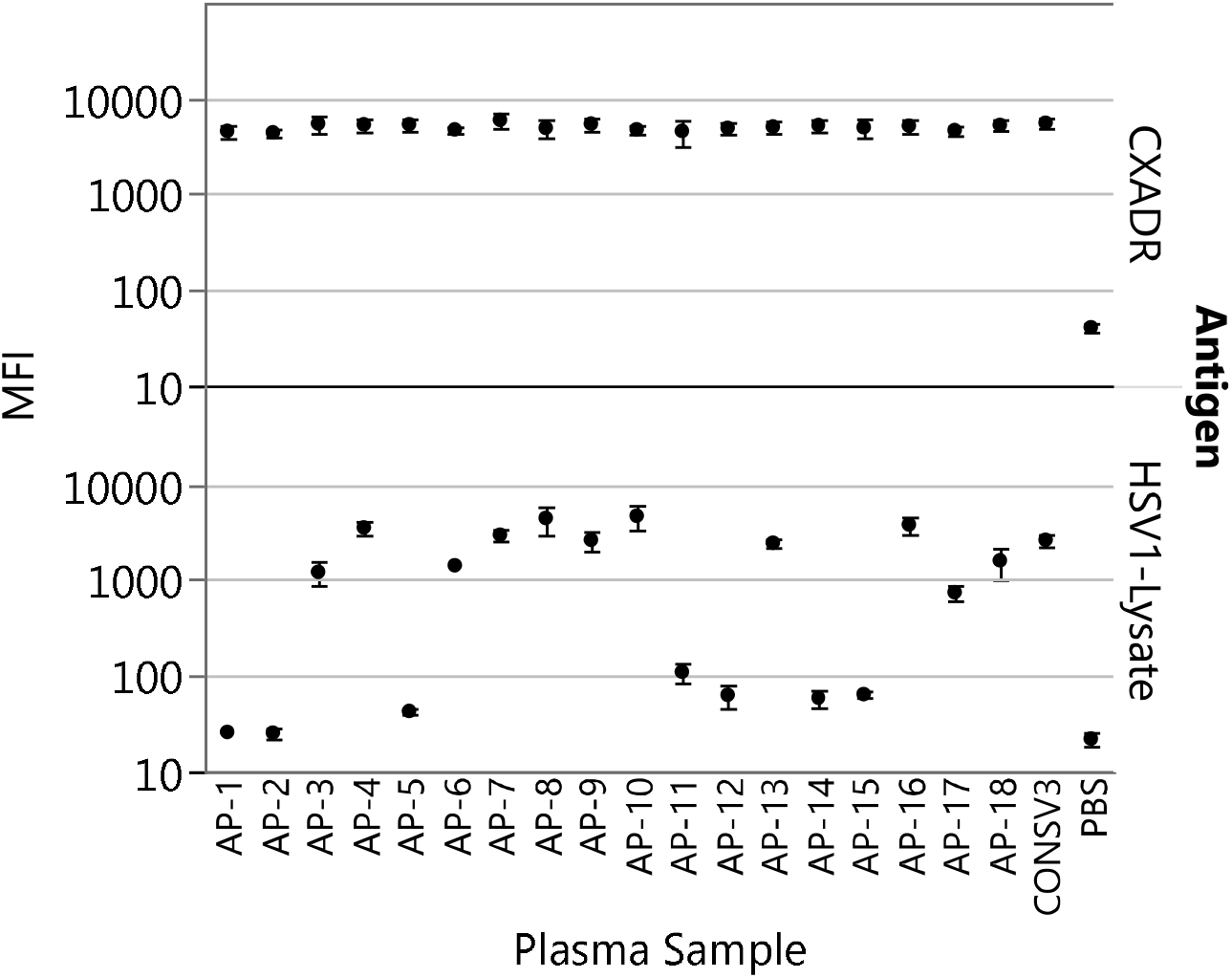
Human plasma (AP-1 to AP-18) diluted 1:1000, CONSV3 positive control diluted 1:1000 and PBS negative control binding to CXADR and HSV1-Lysate coated microspheres.

### Binding deconvolution

The secondary antibody (Goat anti-human IgG/IgA/IgM F(ab’)2 PE from Sigma-Aldrich) that we used for binding of CXADR to human plasma proteins was capable of binding IgA, IgG and IgM Fab domains, so we deconvoluted CXADR binding by measuring human plasma binding with anti-IgA, anti-Mouse IgE, anti-IgM and anti-IgG antibodies (Figure 2), and by binding of human and rabbit IgG1 monoclonal antibodies. IgA, IgE and IgM did not demonstrate binding: plasma binding to CXADR demonstrated no differential signal when read out with anti-IgA, anti-IgM or anti-mouse IgE antibodies (Figure 2), but all of these secondary antibodies demonstrated binding to beads conjugated to human plasma proteins. The IgG1 monoclonal antibodies AUG-3387 expressed in CHO, AUG-3417 expressed in HEK-293, and the rabbit IgG1 transfection control from the Expi293 transfection kit also demonstrated binding to CXADR (Figure 3). Activity against IgD and sub-isotypes IgG2, IgG3, and IgG4 were not assayed.

**Figure 2.**
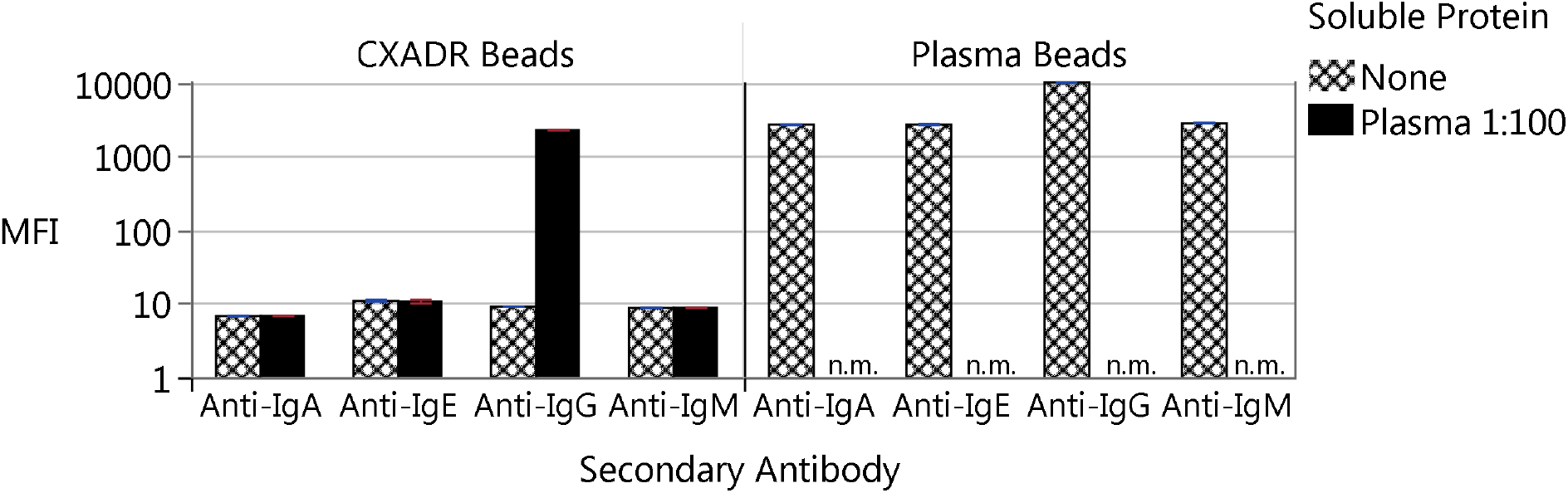
Antibodies from human plasma were tested for binding to CXADR and read out with anti-IgG, anti-IgA, anti-IgM and anti-mouse IgE secondary antibodies. As a positive control, secondary antibodies were also measured for binding directly to beads conjugated to human plasma proteins.

**Figure 3.**
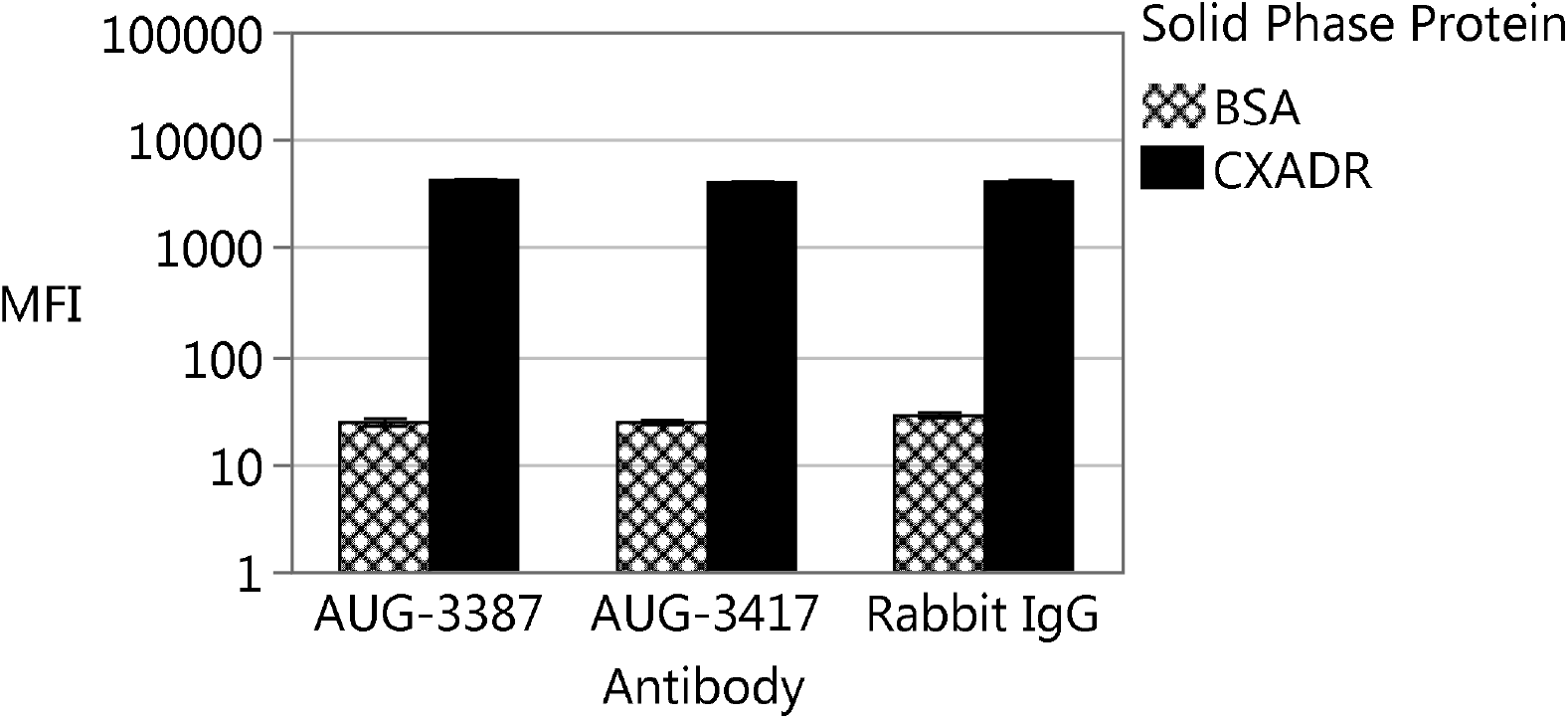
IgG1 monoclonal antibodies AUG-3387, AUG-3417 and the Expi293 rabbit IgG transfection control were measured for binding to BSA and CXADR beads.

### Cross Species Activity

After observing anti-rabbit IgG binding to CXADR, we measured CXADR against other species IgG. We saw no binding of CXADR to mouse, rat, donkey and goat antibodies. Using a rabbit primary antibody and a donkey anti-rabbit secondary antibody we were able to confirm rabbit IgG binding to CXADR (Figure 4).

**Figure 4.**
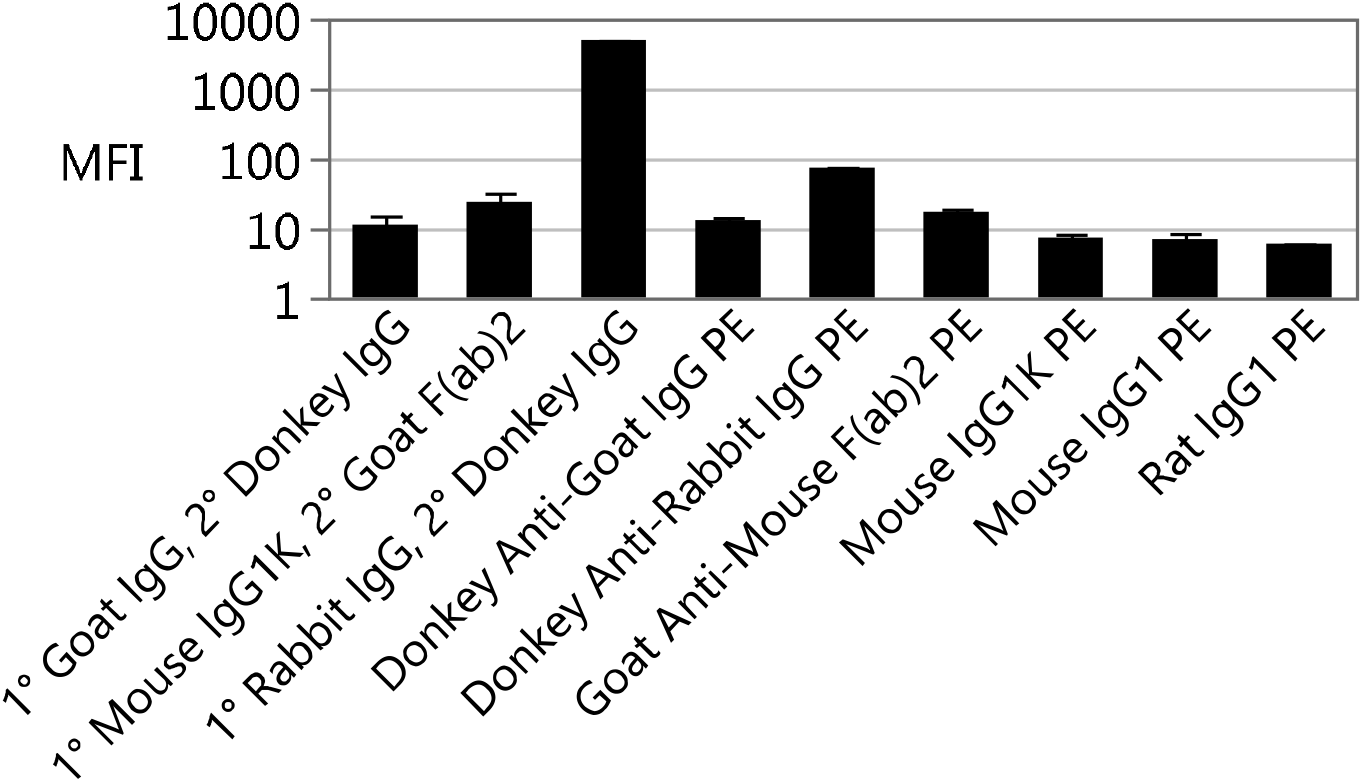
CXADR binding was measured for rabbit IgG, Goat, Donkey, Mouse, and Rat IgG. For the first three conditions, a primary antibody was used with a labelled secondary antibody specific to the primary antibody. The other conditions only included the secondary antibody. All secondary antibodies are labelled with recombinant phycoerythrin.

### Binding affinity

AUG-3387 was titrated from 10 ng/ml to 1 ug/ml against CXADR beads. Binding was dose responsive, and the signal saturated near 300ng/ml. The EC50 determined from a 3 parameter logistic curve (not shown) was 18.6 ng/ml (124pM), and the Kd determined to be 150pM based on fitting the Titration ELISA Curve (Eble, 2018) (Figure 5).

**Figure 5.**
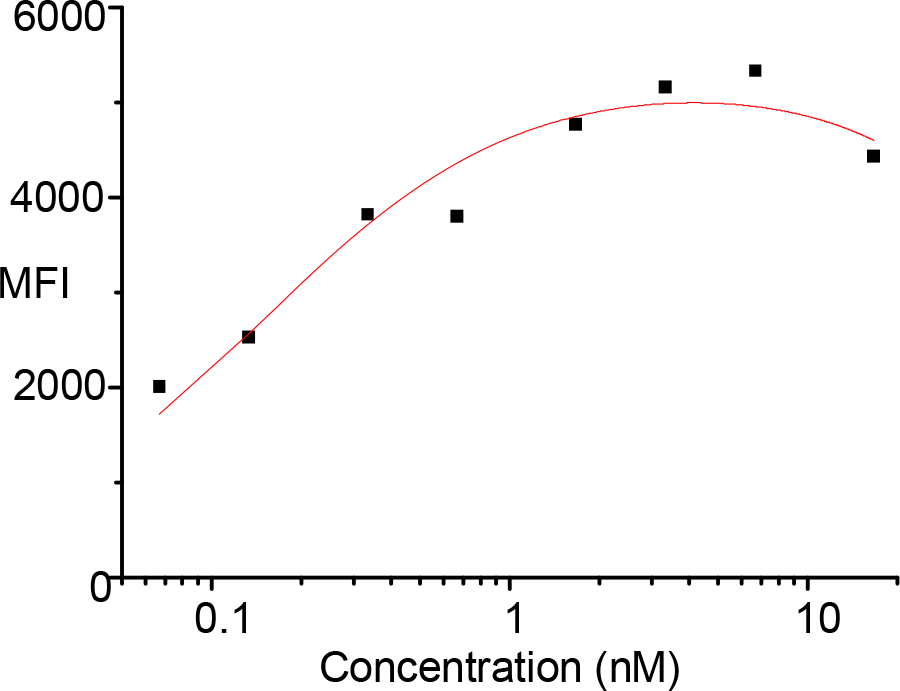
Titration curve of CXADR binding under different concentrations of IgG1. Red line is fit of the Titration ELISA Curve^16^.

### Epitope binning

Epitope binning was performed in two ways: by comparing binding of AUG-3387 to CXADR beads with and without FcBlock reagent (BD, Franklin Lakes, NJ) and by comparing the relative fluorescence with and without AUG-3387 after staining with anti-Fc secondary antibodies versus Anti-Fab antibodies. Up to a 5 fold reduction in signal was observed by including FcBlock reagent before/during incubation with 1ug/ml AUG-3387 (Figure 6). No change in fluorescence intensity was detected upon secondary staining with anti-Fc antibodies after binding AUG-3387 to CXADR beads, but anti-Fc antibodies were clearly detectable when staining beads containing human plasma proteins, indicating that anti-Fc antibodies were unable to bind AUG-3387 in complex with CXADR (Figure 7). Anti-Fab antibodies on the contrary demonstrated a greater than 100X increase in signal. Confirming these results and consistent with IgG specificity, CXADR did not bind AUG-3387 when expressed as an ScFv (Figure S1).

**Figure 6.**
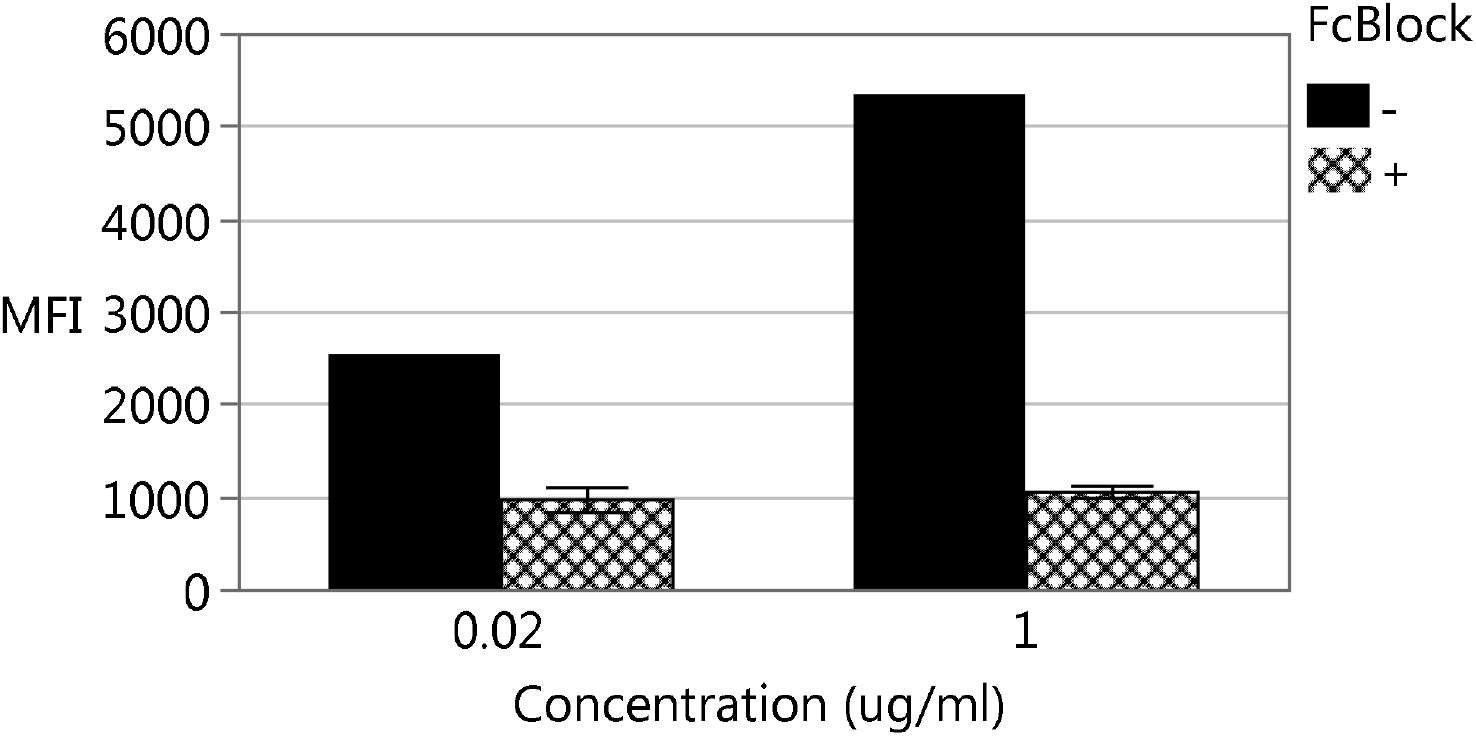
Binding of AUG-3387 at two concentrations to CXADR with and without FcBlock.

**Figure 7.**
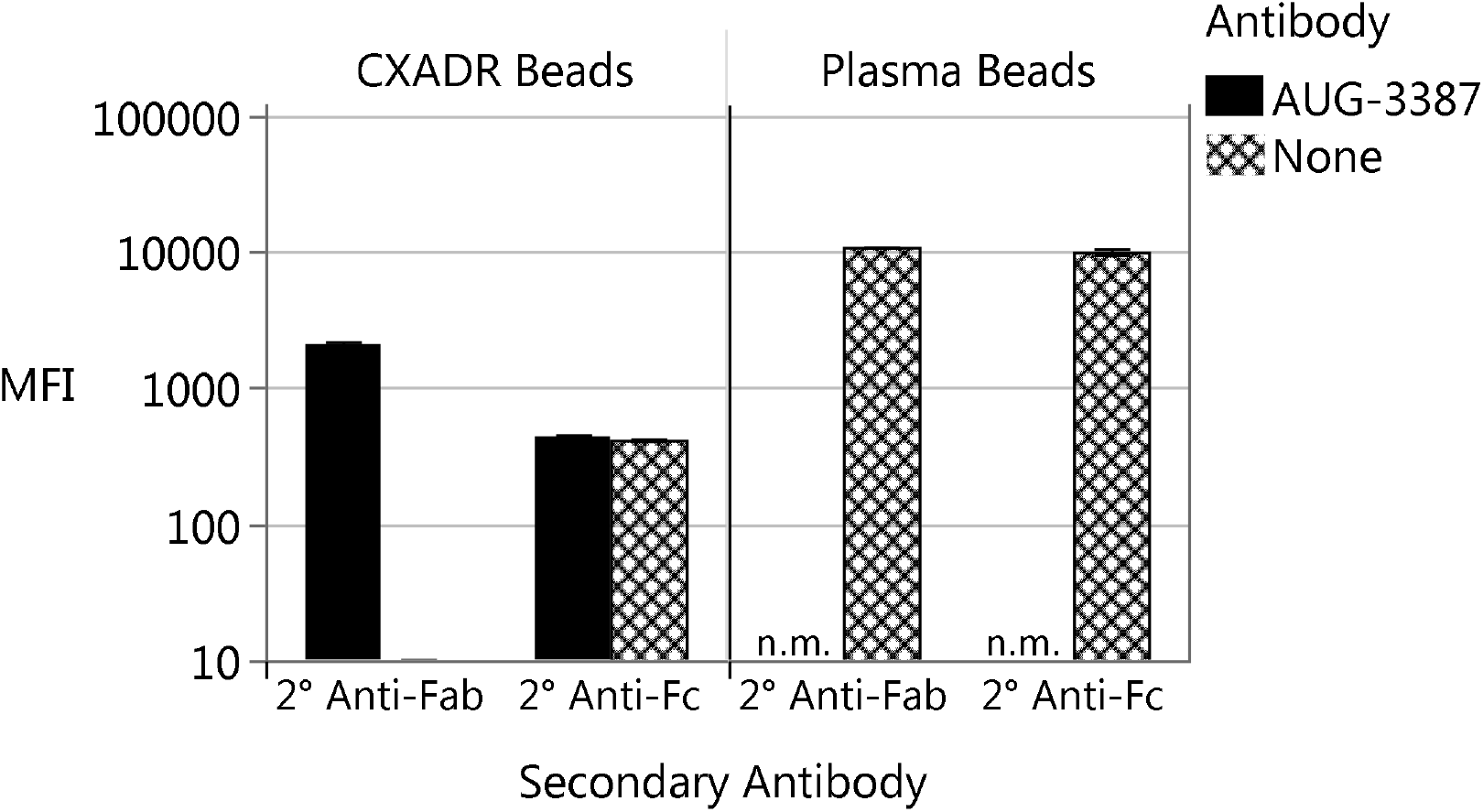
Binding of AUG-3387 to CXADR as read out with anti-Fab and anti-Fc antibodies (left). Secondary antibodies were also measured for binding to human plasma proteins as a positive control (right).

## Conclusion and Discussion

CXADR binds human IgG and not IgA or IgM and likely not IgE. FcBlock inhibits CXADR-IgG binding and CXADR inhibits readout with anti-Fc secondary antibodies indicating CXADR binds the Fc portion of human IgG. Based on titration ELISA data, we estimate the binding affinity to be approximately 1nM. CXADR with its two surface exposed Ig domains has structural homology to Fc receptors and Fc receptor like proteins ^17^. We conclude that CXADR has a previously unidentified role as a high affinity human Fc gamma receptor.

FcR’s are a crucial class of receptors in human biology. FcR’s are expressed in immune tissues, barrier epithelium and the human kidney and are key components in numerous processes such as antibody mediated endocytosis, ADCC, B cell expansion and differentiation, and antibody recycling^18,19^. FcR’s are crucial for fighting infections, B cell maturation, and maintaining peripheral immune tolerance^20–23^. Human diseases with known roles for FcR’s include RA, APS, SLE, Sjogren’s, anaphylaxis, Hashimoto’s thyroiditis, allergies, cancer and nearly every type of infection^22,24^. FcR’s have a dominant role in determining the half-life of circulating antibodies, directly impacting the pharmacodynamics of full chain monoclonal antibody therapeutics and Fc fusion biologics ^25–34^.

While CXADR expression is relatively low, CXADR is expressed in most human tissues. Based on data from the Human Protein Atlas^35–39^, we estimate that CXADR represents approximately 0.5 percent of all FcR in the body, similar in magnitude to FcγRI (Figure S3). It remains an open question if CXADR is bound to IgG in vivo and in what ways that would affect human physiology. Given the high concentration of IgG in the body, the data provided herein suggests CXADR would be considered a “high affinity FcR” and it is possible that CXADR is a substantial contributor to the total fraction of Ig bound to FcR.

This new activity for CXADR has implications for future experiments and previous results. Perhaps most poignantly, given that previously executed experiments do not account for the Fc binding nature of CXADR and its wide ranging tissue distribution, it is possible that some immunohistochemistry and cytometry results may need to be reconsidered if they utilized rabbit or human IgG in staining protocols. Regarding the physiological role of CXADR as an FcR, Ig bound to CXADR may be involved in immune surveillance at tight junctions via direct inhibition of antigens, antigen presentation, immune signaling, or cell trafficking. For instance, CXADR could serve as an attachment point for cells with surface bound Ig during extravasation into infected tissues. Interactions between CXADR and IgG could lead to changes in barrier permeability at the blood-testis barrier, or blood brain-barrier which would have implications for auto-immune infertility and Long Covid^40^. CXADR could serve as a sink for circulating IgG, and be responsible for immune downregulation toward homeostasis. Receptor-receptor interactions involving CXADR could be inhibited by IgG and IgG could induce CXADR crosslinking in cis-acting or trans-acting capacity. Antigen bound Ig bound to CXADR may be internalized and re-presented via Type I MHC, drawing pro-inflammatory cells to the site of an infection via CD8 T cell responses. Since nervous system and cardiac tissues have cells in which CXADR is expressed in the absence of any other known FcR ^35–39^ (Figure S4), these processes may have implications for cardiac and neural inflammation.

Interactions between antibodies, antigens and CXADR could be implicated in the development of several diseases relating to immune dysregulation. For instance, it is possible that CXADR is responsible for antibody dependent enhancement seen with some viral infections. Given that CXADR is a receptor for several viruses, even “neutralized” virus particles may bind target cells via antibody-FcR interactions, indicating the evolutionary importance of these entry receptors from both a host and virus perspective. Non-neutralizing antibodies bound to SARS-CoV-2, could increase infection of CXADR expressing cells via ADE or crosslink CXADR and ACE2 potentially leading to aberrant signaling with unpredictable effects. Alternatively, antigens bound to CXADR through antibodies could be internalized and cross-presented through Type I MHC, potentially initiating a disastrous cytotoxic T cell response to critical cells, e.g., oligodendrocytes, HCRT neurons, beta islet cells, or cardiomyocytes implicated in diseases such as multiple sclerosis^41^, narcolepsy^42^, Type 1 diabetes^14,43,44^ and Covid-19 induced cardiomyopathy^8,45,46^, respectively. This type of antigen cross presentation could also have implications for sepsis in response to bacterial antigens^47,48^, hemorrhage from viral infection (e.g. Dengue Hemorrhagic Fever)^49–51^, or neuroinflammation in response to HSV1 associated Alzheimer’s Disease^52^. While these are purely hypothetical mechanisms for discussion purposes, given the many tissues that express CXADR of various isoforms and the criticality of Ig related immune processes in disease, it is an area worth further investigation.

## Methods

### Human Plasma and Antibody Expression

Rabbit IgG was expressed in ThermoFisher (Waltham, MA) Expi293 cells in a 24 well deep well block according to the manufacturer’s instructions.

AUG-3387 IgG1^15^ was expressed in CHO cells by Catalant (Somerset, NJ), and AUG-3417 IgG1^15^ was expressed in HEK293 cells by Absolute Antibody (Oxford, UK). AUG-3387 ScFv (a.k.a. AUG-3705) ^15^ with N Terminal 6xHis, Myc and V5 peptide tags was expressed in E. coli, and purified via anion exchange FPLC.

Human blood was obtained from study volunteers under informed consent. 10ml of human blood was obtained in ACD tubes. Plasma was separated from cells and platelets after centrifugation at 1000g and stored at -80C until needed. All samples from human participants were collected after providing their written consent according to an IRB approved protocol (Quorum IRB 32963). All research was performed in accordance with the IRB protocol, the Declaration of Helsinki, and local laws and regulations.

### Bioconjugation

A solution of forty micrograms of protein (CXADR or human plasma) was loaded onto a 0.5ml Zeba 7k MWCO spin filter and centrifuged at 1500g, then washed three times with 300ul of PBS centrifugation at 1500g. An aliquot of 20 micrograms of the washed protein was conjugated to 5 million Luminex magnetic microspheres according to recommendations in the Luminex Bio Conjugation Kit. Bead solutions were blocked with 0.1% StabilGuard for 30 minutes, decanted, and resuspended in 0.1% PBS-Tween20. Beads were quality controlled by performing a NanoOrange assay directly on conjugated protein indicating at least 1ug protein coupled per 1 million beads.

### Luminex reactions

Luminex magnetic microspheres conjugated to CXADR ectodomain or human plasma were decanted and resuspended in PBS BSA buffer, and a solution of 10ul containing 500-1000 beads was added to assay plates. Antibodies, or human plasma were diluted in 90ul PBS-BSA and incubated with the beads for 60 minutes. Plasma was diluted at 1:100 or 1:1000 final in assay buffer, and antibodies were diluted at various increments in the 10ng/ml to 10ug/ml range of final concentrations. Following a 60 minute incubation, bead solutions were decanted and washed twice with 80ul of PBS-Tween buffer. Secondary antibodies conjugated to recombinant phycoerythrin were diluted in 95ul PBS BSA buffer at a final concentration of 1ug/ml. Following a 30 minute incubation with secondary antibody, bead solutions were decanted and washed twice with 80ul of PBS-Tween (0.1% v/v) buffer, resuspended in 80ul of PBS-BSA buffer and then assayed for median fluorescent intensity on a BioPlex 200 instrument.

### Epitope Binning

FcBlock inhibition of CXADR-antibody binding was assayed by addition of 1ul of FcBlock reagent to the bead solution before addition of AUG-3387 IgG1. FcBlock was added at 9% V/V and incubated at room temperature for 10 minutes before adding AUG-3387 at which point it was present at 1% V/V concentration. A positive control was performed without addition of FcBlock in the same experiment.

## Data Analysis

Curve fits and plots were performed in JMP and OriginPro. EC50 was determined by fitting concentration (ug/ml) and MFI to a 3 parameter logistic equation and extracting the inflection point. Kd was determined by fitting concentration (nM) and MFI to the “Titration ELISA Curve” Equation 9 from Eble (Eble, 2018) :

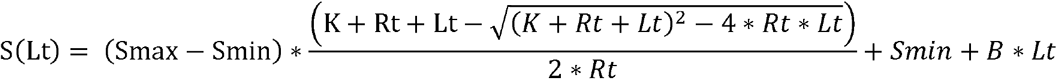

## Materials

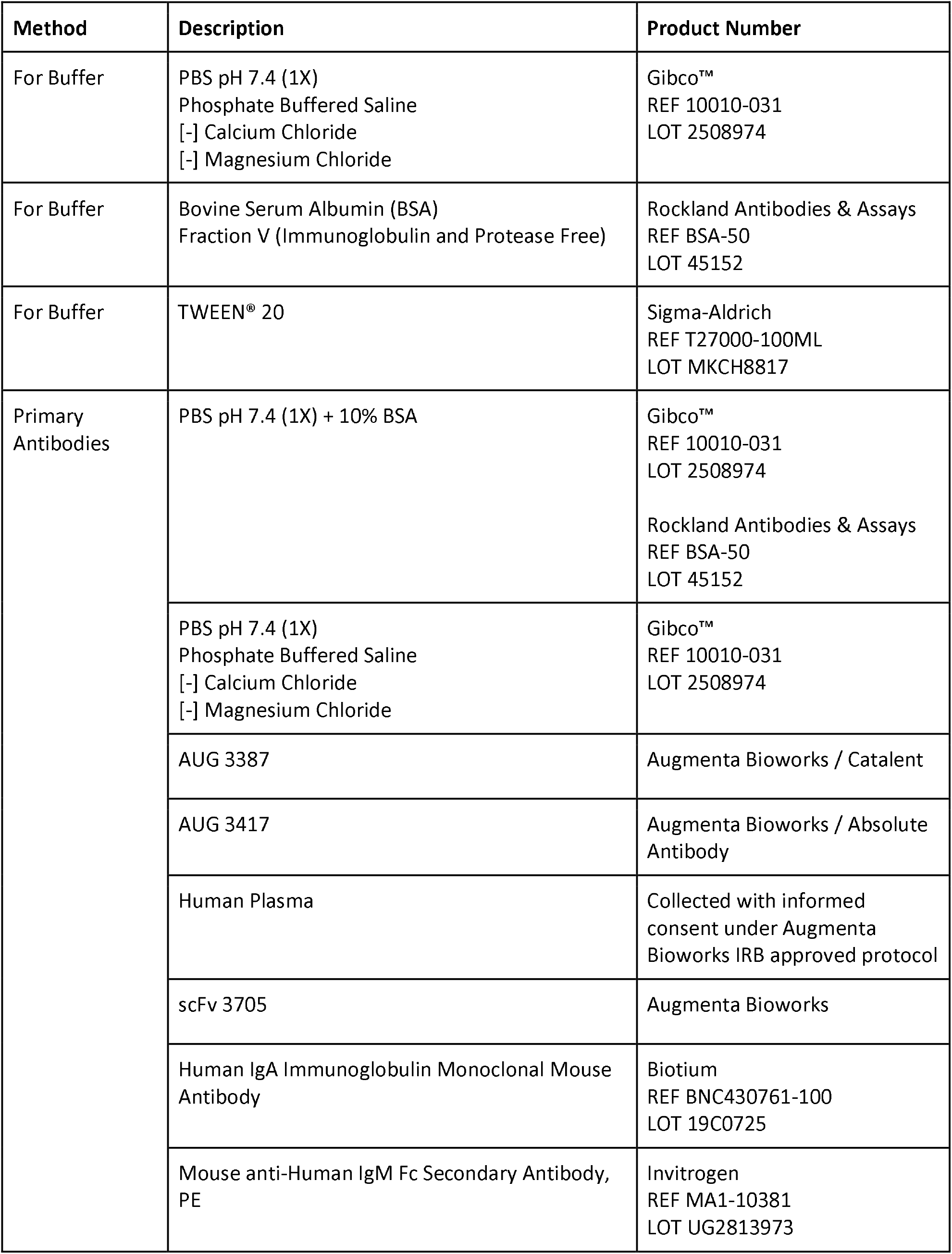
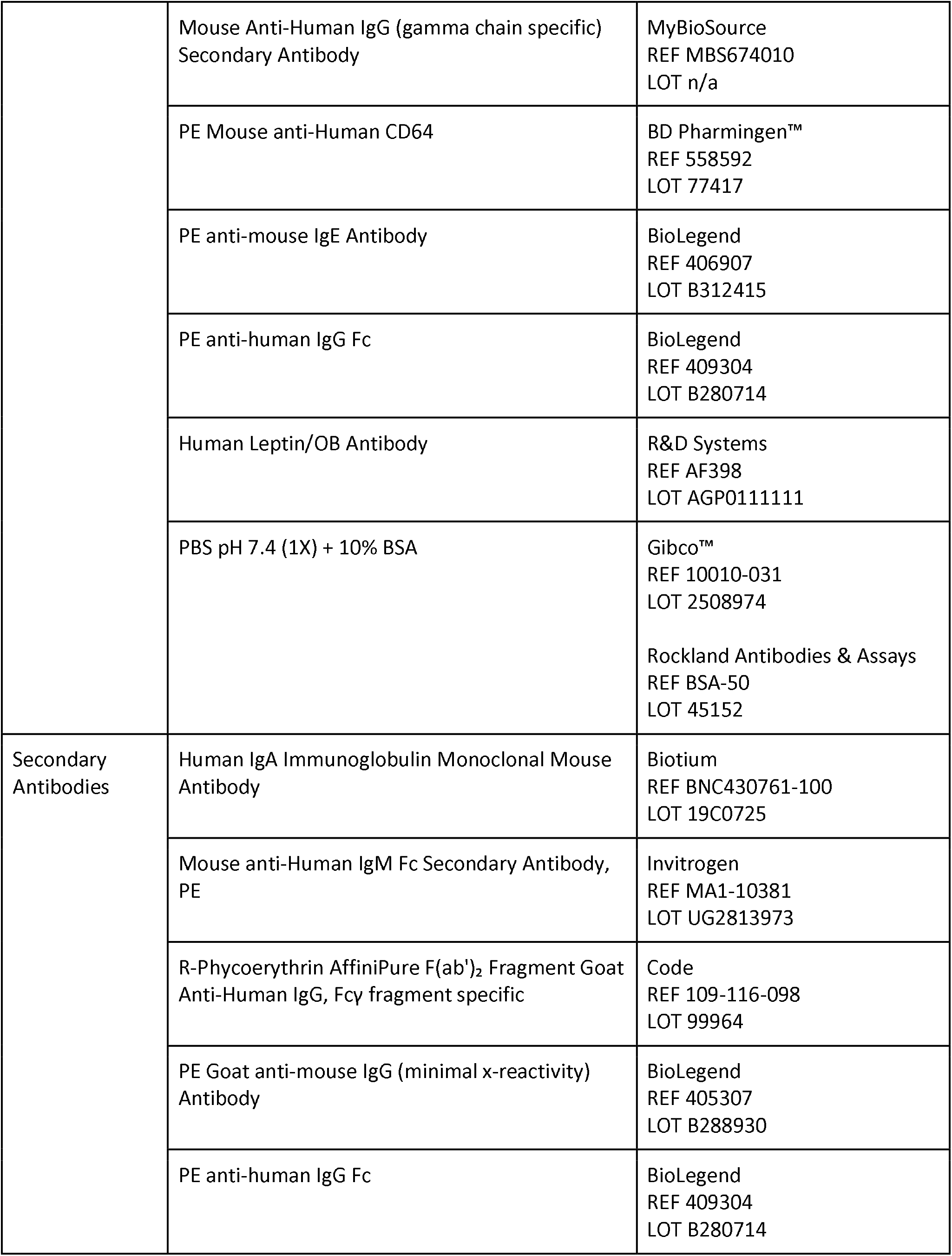
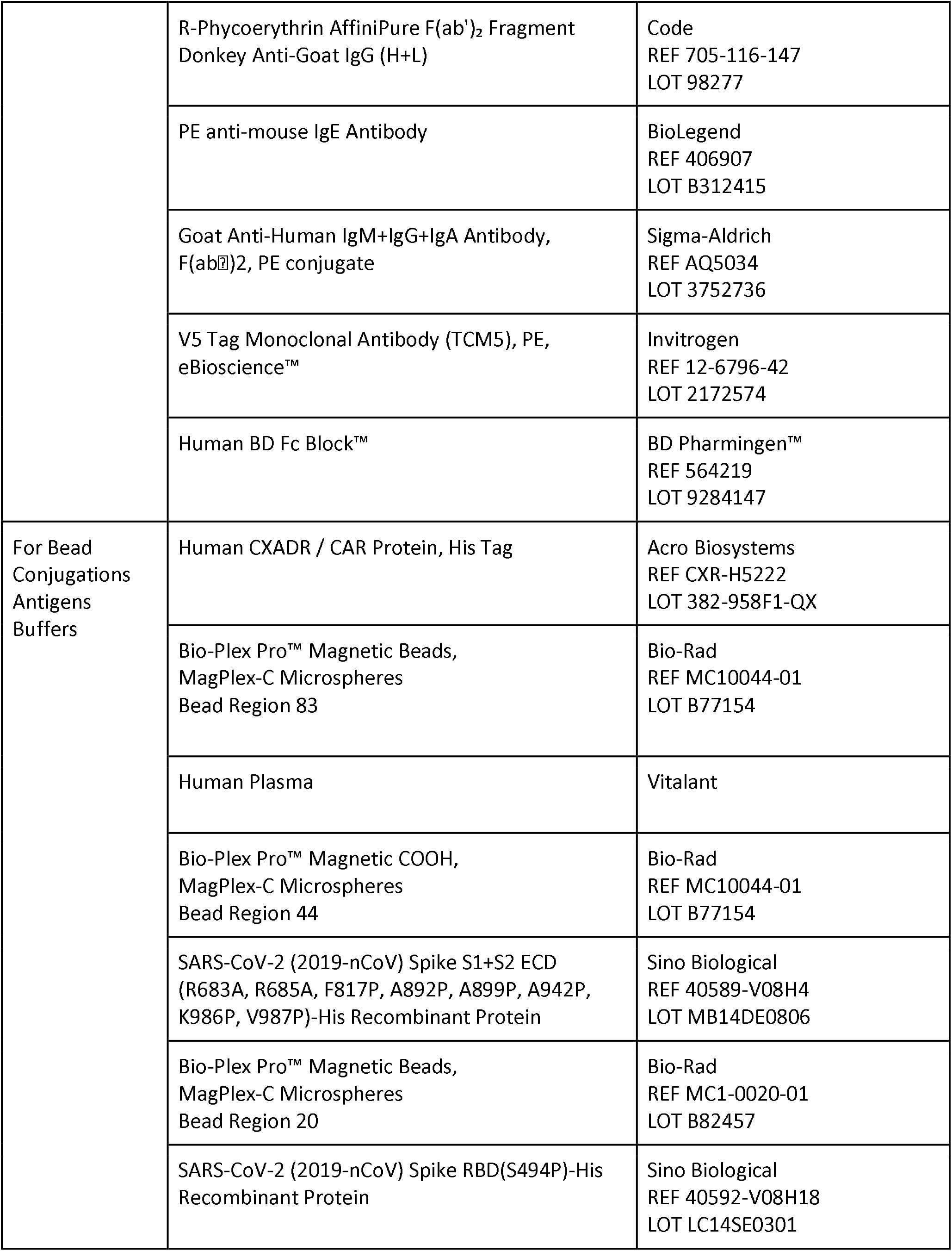
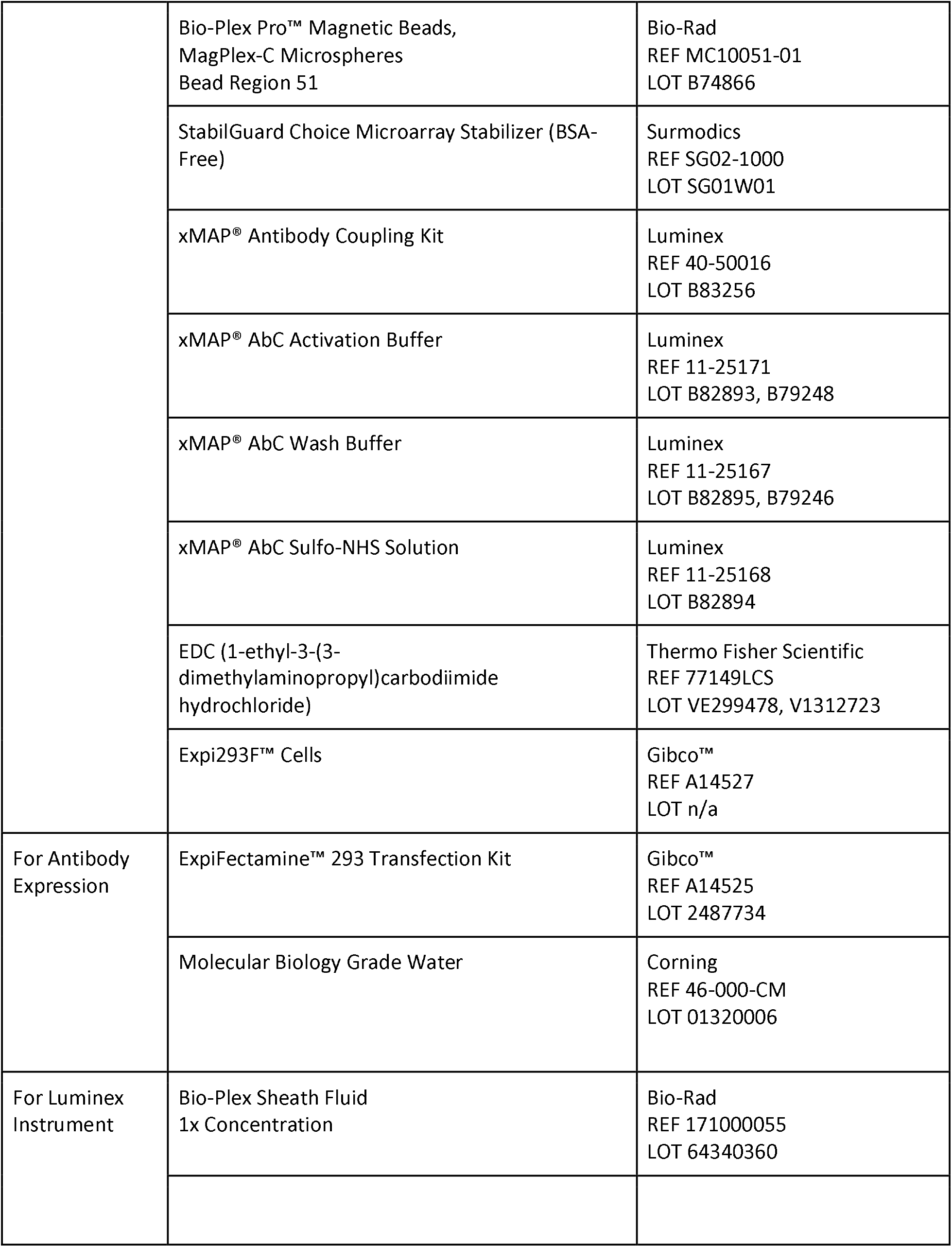

## Acknowledgements

The authors would like to express their gratitude for the blood sample donations from study participants and the ME/CFS serological data generated in collaboration with the Amar Foundation and the Maureen Hanson Lab of Cornell University. We would also like to thank Steven P. Henry, Payam Shahi, and Marco A. Mena for research materials, helpful discussions and logistical support.

## Data Availability

The datasets used and/or analyzed during the current study available from the corresponding author on reasonable request.

