## Supplemental data for "CXADR is a Human IgG Fc Receptor"

Supplementary Data

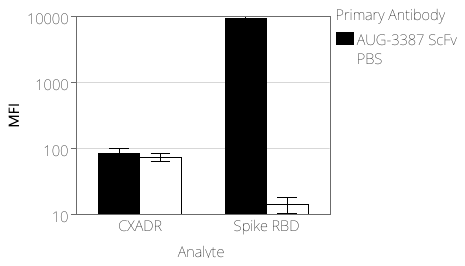

Figure S1.

AUG-3387 ScFv (AUG-3705) binding to CXADR and Spike RBD beads, read out with anti-V5 secondary antibody.

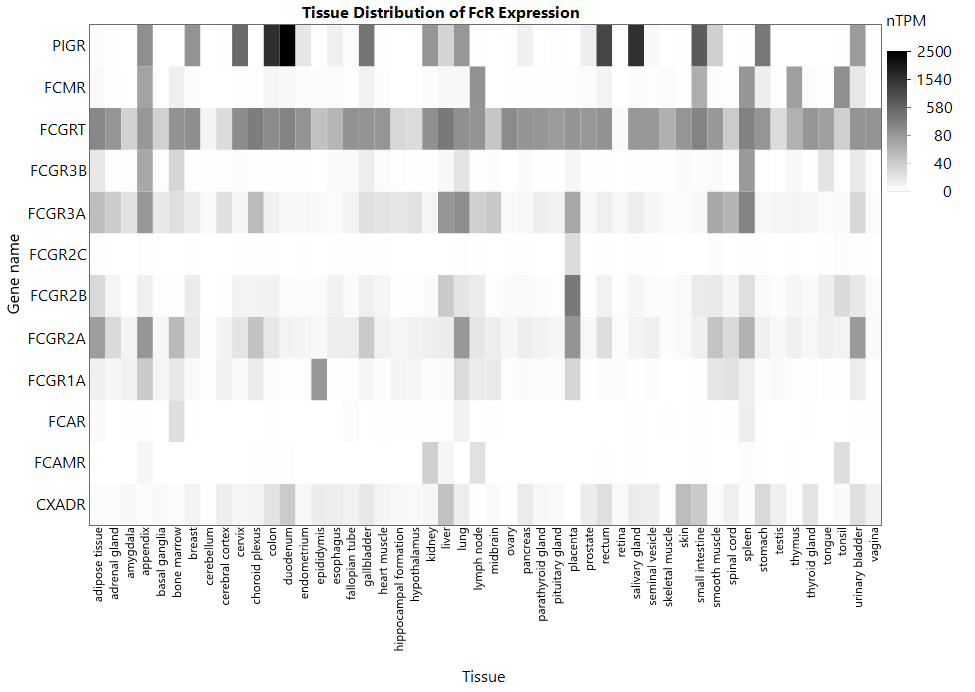

Figure S2. FcR and CXADR expression from The Human Protein Atlas version 23.0 ^39^

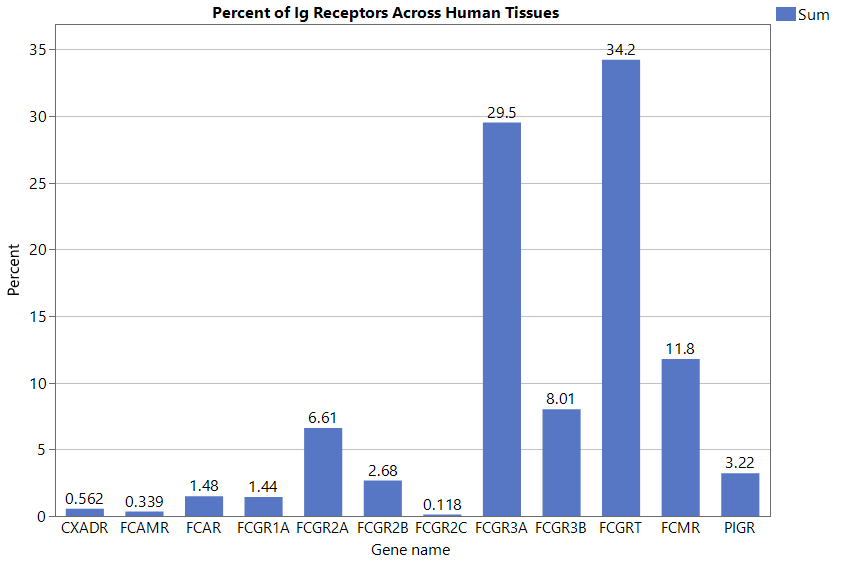

Figure S3. Approximate aggregate whole-body FcR and CXADR expression as a percent of total FcR generated by merging data from the Human Protein Atlas version 23.0 ^39^ and cell count data from ^53^, excluding counts associated with duodenum, endometrium, ovary, and fallopian tubes.

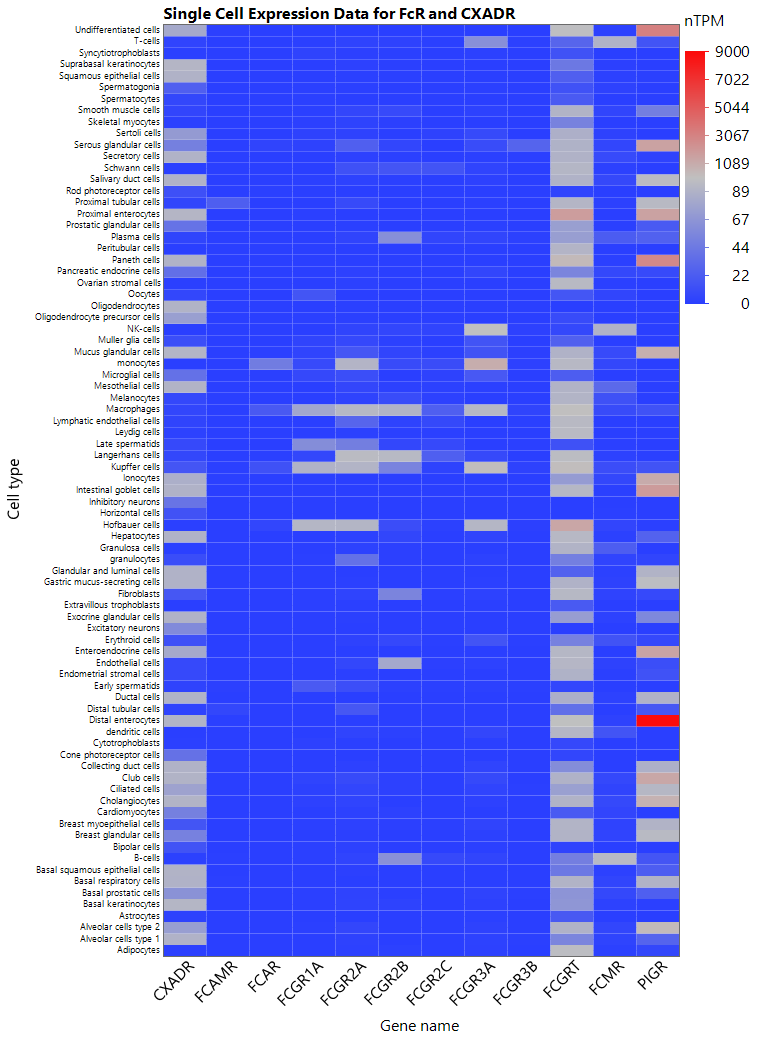

Figure S4. Single cell expression data for CXADR and FcR ^35^.

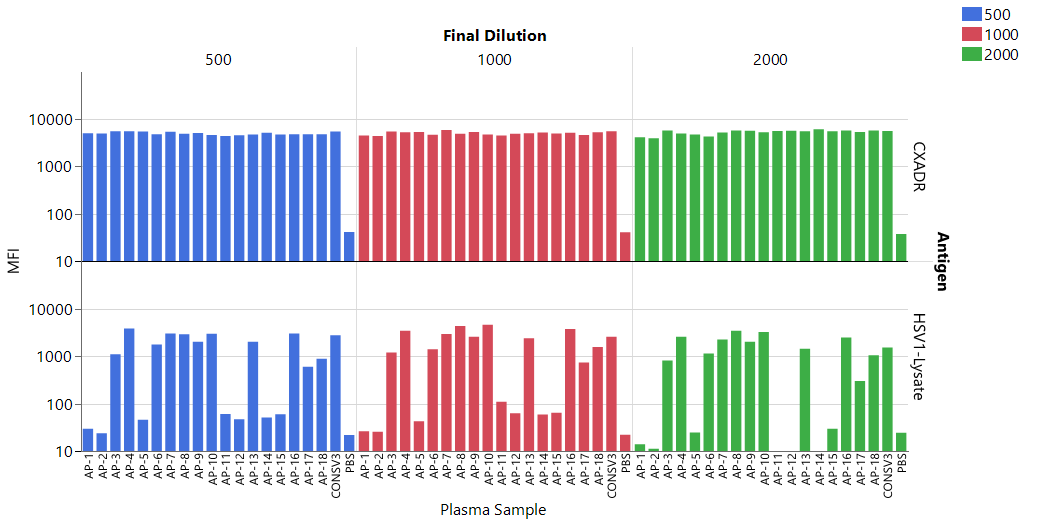

Figure S5. Plasma titrated from 1:500 to 1:2000 dilutions bound to CXADR and HSV-1 lysate microspheres.
